# Single-cell analysis of *Schistosoma mansoni* reveals a conserved genetic program controlling germline stem cell fate

**DOI:** 10.1101/2020.07.06.190033

**Authors:** Pengyang Li, Dania Nanes Sarfati, Yuan Xue, Xi Yu, Alexander J. Tarashansky, Stephen R. Quake, Bo Wang

## Abstract

Schistosomes are parasitic flatworms causing one of the most prevalent infectious diseases from which millions of people are currently suffering. Their germline outputs many fertilized eggs, which are both the transmissible agents and the cause of the infection-associated pathology. Given its significance, the schistosome germline has been a research focus for more than a century. Nonetheless, little is known about the molecular mechanisms that regulate its development. Here, we construct a transcriptomic cell type atlas of juvenile schistosomes. This allows us to capture germline stem cells (GSCs) during *de novo* gonadal development. We identify a genetic program that balances the fate of GSC between proliferation and differentiation. This program is controlled by *onecut*, a homeobox transcription factor, and *boule*, an mRNA binding protein. Evaluating this genetic program in schistosome’s free-living evolutionary cousin, the planarian, shows that this germline-specific regulatory program is conserved but its function has changed significantly during evolution.

## Introduction

Schistosomes are parasitic flatworms that cause one of the most prevalent but neglected infectious diseases, schistosomiasis^1^. With over 250 million people infected worldwide and a further 800 million at risk of infection, schistosomiasis imposes a global socioeconomic burden comparable to that of tuberculosis, HIV/AIDS, and malaria^1–3^. The disease transmission requires the passage of parasites through two hosts during its life cycle, a molluscan intermediate host and a mammalian definitive host (e.g., human). This complex life cycle requires the deployment of multiple specialized body plans in order to infect and reproduce within each host. These transitions are enabled by a population of stem cells that undergo several waves of proliferation and differentiation^4–6^.

The schistosome life cycle begins with the parasite egg being excreted from the mammalian host into fresh water, which hatches into a free-swimming larva that serves as a vehicle for approximately a dozen stem cells to penetrate a snail host and transform into a sporocyst^4,5,7,8^. Stem cells in the sporocyst can undergo either self-renewal or enter embryogenesis, producing more sporocysts or massive numbers of infectious progeny (called cercariae), respectively^4,5,8–10^. Cercariae then emerge from the snail back into water, burrow through the skin of a mammalian host, migrate to species-specific niches in the host vasculature and transform into juvenile parasites. This transition initiates the sexual portion of the life cycle. Stem cells packed in cercariae proliferate to initiate the growth of juveniles and eventually build sexual reproductive organs *de novo*, through a developmental program that appears to be modulated by host hormonal and immune cues^5,11,12^. Sexually mature male and female worms pair to produce fertilized eggs, which are excreted to initiate the next life cycle. It is worth noting that the eggs, not the worms, cause pathology as they become trapped in the host tissues and induce extensive granulomatous inflammation^13^.

We recently conducted a single-cell transcriptomic study comparing stem cells from *Schistosoma mansoni* asexual (intramolluscan) and sexual (intramammalian) stages^5^. Among the stem cells that are carried by cercariae and transmitted from snail to mammal, only a subset of them commits to the germline fate. These cells initially express a novel schistosome factor, *eledh* (*Sm-eled*, for brevity the prefix ‘Sm’ will be omitted from gene names in the rest of the paper wherever species origin is unambiguous), and during germline specification activate one of the schistosome homologs of *nanos* (*nanos-1*), an RNA binding protein conserved across metazoans^5,14–18^. Unlike most animals that segregate their germline from soma during embryonic development^15,19,20^, schistosomes specify their germ cells from a somatic cell lineage at the onset of juvenile development.

While our previous work has identified the origin of the schistosome germline, it remains unclear which molecular regulatory program(s) underlines the separation of somatic and germ cell lineages. In contrast to their distinct cellular fates, somatic and germline stem cells share strikingly similar gene expression signatures, including genes involved in regulating cell cycle, a set of conserved RNA binding proteins that are often associated with stem cell multipotency^4–6^, and transcription factors such as nuclear factor Y subunits^18^. Disrupting these key regulators leads to defects in stem cell maintenance and proliferation in both somatic and germline lineages^4,18^. The molecular distinction between somatic and germline stem cells is thought to be similarly blurry in other flatworms, such as schistosome’s free-living evolutionary cousin, the planarian^19,21^. Planarian germline stem cells resemble their somatic stem cells both in terms of morphology and molecular signatures, and can even contribute to somatic tissues during tissue regeneration after injury^20–24^.

Here, we use single-cell RNA sequencing (scRNAseq) to construct a transcriptomic cell type atlas of juvenile *S. mansoni* to isolate and characterize the germline stem cells (GSCs). We focus on this life-cycle stage because juvenile schistosomes contain abundant stem cells, including cells in the transition between somatic and germline fates^5^. Enabled by the self-assembling manifolds (SAM) algorithm^25^, a method that excels in identifying subtle transcriptional differences between otherwise similar groups of cells, we succeeded in capturing the GSCs and identifying their transcriptional signatures. Through RNA interference (RNAi) mediated gene knockdown, we evaluated the function of GSC-specific transcripts and found a schistosome homolog of *onecut* homeobox transcription factor, *onecut-1*, to be the key regulator that balances the proliferation and differentiation of GSCs (in particularly, male GSCs, or spermatogonial stem cells). *onecut-1* functions through complex epistatic interactions with several other genes, including *eled, nanos* and *boule*. As *nanos* and *boule* are broadly conserved germline developmental regulators^14–18,26–30^, and *onecut* expression has also been detected previously in mammalian testes^31–34^, we posit that this gene set may represent a conserved germline specific regulatory program. Consistent with this idea, we found that *onecut*, along with *boule* and *nanos*, are indeed essential in the male germline of planarian *Schmidtea mediterranea*, though their specific function has changed. While *onecut* protein family has been extensively studied across animals (e.g., fly and mouse)^32–37^, its role in regulating germline development has been previously unknown.

## Results

### scRNAseq identifies GSC-specific transcriptional signatures

We sequenced all body cells in juvenile schistosomes using Smart-seq2 to improve the detection of lowly expressed genes including transcription factors (TFs)^38–40^. We collected 10,945 single-cell transcriptomes, of which 7,657 passed the initial quality control (see **Methods**). **Fig. 1a** shows that SAM analysis divided these cells into several major clusters, but only one population expresses *ago2-1*, an argonaute homolog that is ubiquitously and specifically expressed in all schistosome stem cells^4–6,25^. Performing SAM analysis on *ago2-1^+^* cells (1,464 cells) separately revealed nine subpopulations (**Fig. 1b**), four (*μ*, *δ*′, *ε_α_* and *ε_β_*) of which we reported earlier^5,25^. The five newly found populations include one that contains exclusively cells expressing *nanos-1^+^*, a previously characterized GSC marker^5,18^. The other four contain cells expressing genes often associated with differentiated tissues (**Supplementary Table 1**), for example, *troponin* (a common muscle gene^25,41^), *complexin* (*cpx*, a neural gene), *cathepsin B endopeptidase* (*cb2*, a gene expressed in intestinal and parenchymal cells^42,43^), and *tetraspanin-2* (*tsp-2*) co-expressed with *Sm25* (both are known markers for schistosome epidermal lineage^44^). Since these populations still express common stem cell markers (e.g., *ago2-1, cyclin B, pcna, polo-like kinase*, histone *h2a* and *h2b*) and are actively dividing (**Supplementary Fig. 1**), they are likely tissue type-specific progenitors, though further functional validation would be required to prove their respective fates.

**Figure 1:**
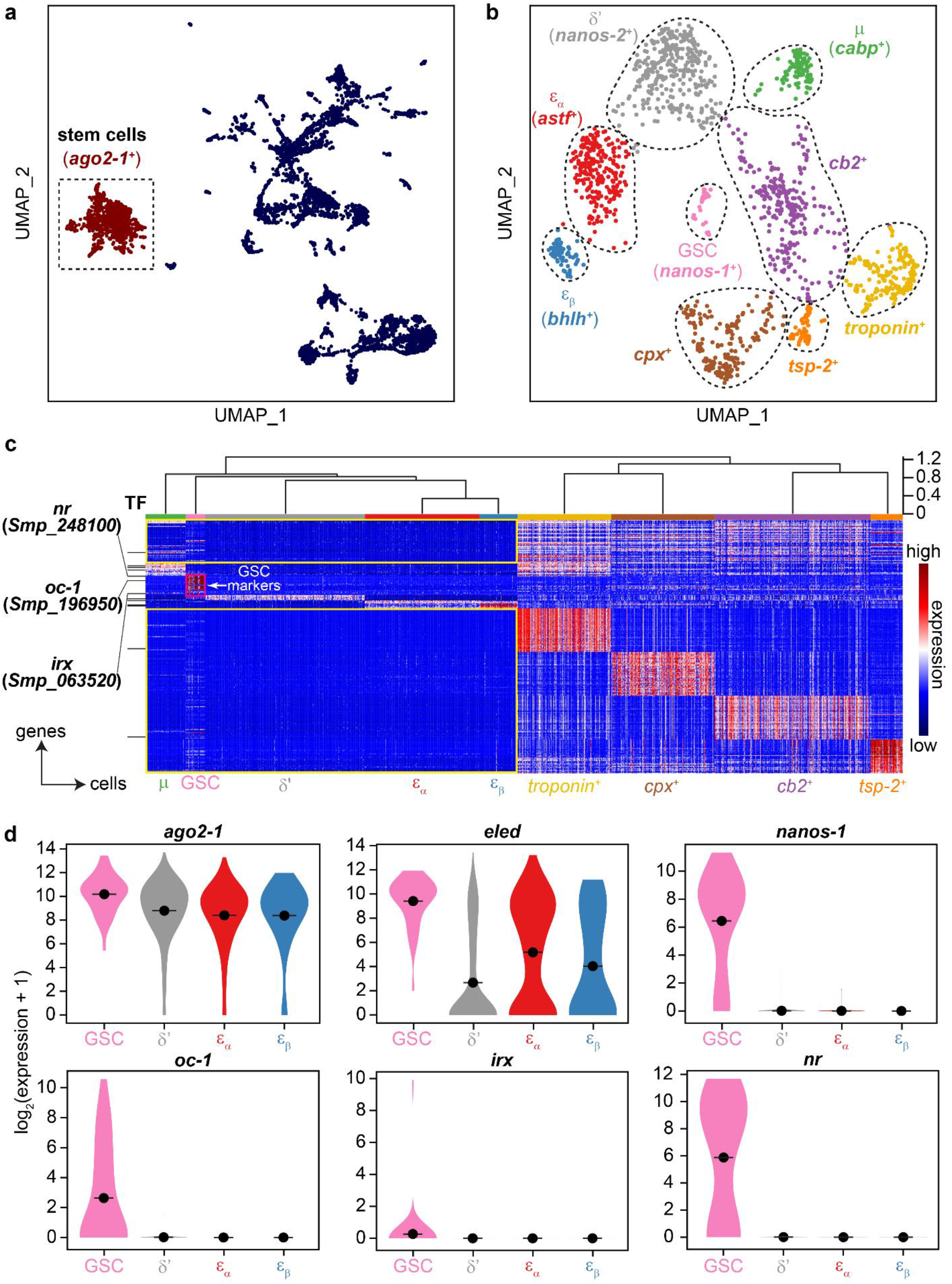
scRNA-seq identifies schistosome stem cell subpopulations and GSC-specific transcriptional signatures. UMAP projections showing scRNA-seq data of (a) all cells and (b) *ago2-1^+^* stem cells only. The manifolds are reconstructed by SAM^25^. In (a), *ago2-1^+^* stem cells are highlighted in red. In (b) cells are color-coded by subpopulation annotations determined by marker gene expression. (c) Scaled expression heatmap showing population specific marker gene expression. The expression for each gene was standardized to have zero mean and unit variance. Top: the subpopulations are grouped through hierarchical clustering in principal component (PC) space, the height of the branches represents correlation distance between populations. Left: annotated transcription factors (GO: 0003677 DNA binding and GO: 0003700 DNA-binding transcription factor activity) are indicated by black lines. Yellow boxes: genes that lack expression in the presumptive multipotent group compared to the progenitor group. Red box: genes specifically expressed in GSCs. (d) Violin plots showing distributions of expression levels for *ago2-1* (an ubiquitous stem cell marker), *eled* (enriched in GSCs), *nanos-1* (a previously known GSC marker), and the three identified GSC-specific TFs, *oc-1, irx* and *nr* in GSC, *δ*′-, *ε_α_*- and *ε_β_*-cells. Dot: mean expression within each population.

Hierarchical clustering of the cell populations (see **Methods**) divides them into two major groups, one grouping *μ*-, *δ*′-, *ε_α_*- and *ε_β_*-cells together with GSCs, and the other containing the four putative progenitor populations (**Fig. 1c**). Unlike progenitor populations that express a large number of population-specific markers, *μ*-, *δ*′-, *ε_α_*-, *ε_β_*-cells and GSCs only express a small number of population-specific genes beyond the common stem cell markers (**Supplementary Table 1**). Instead, this group is separated from the progenitor group primarily based on the lack of marker gene expression (yellow boxes in **Fig. 1c**), consistent with the idea that multipotent stem cells repress genes that are involved in cell differentiation^45^. Among the presumptive multipotent populations, we identified a total of 22 genes that are specifically expressed in GSCs with respect to all other cells in our dataset (red box in **Fig. 1c**). This gene set includes three TFs (**Fig. 1d**): *onecut-1* (*oc-1*, Smp_196950, **Supplementary Fig. 2**), a homolog of iroquois homeobox TF (*irx*, Smp_063520), and a putative nuclear hormone receptor (*nr*, Smp_248100), all of which are expressed in subsets of GSCs.

### *oc-1* suppresses proliferation and promotes differentiation of male GSCs

Whole-mount in situ hybridization (WISH) revealed distinct expression patterns for the three identified TFs (**Fig. 2a**). *oc-1* expression is specific to primordial testes. *irx* is expressed in both testes and ovary. While *nr* is not expressed in male or female gonadal primordia, it is co-expressed with *oc-1* in primordial vitellaria of female parasites, consistent with the presence of *nanos-1^+^* cells in this yolk cell-producing somatic reproductive organ^46^. Fluorescence in situ hybridization (FISH) experiments also confirmed that *oc-1* is expressed in a subset of *nanos-1^+^* spermatogonial stem cells (**Supplementary Fig. 3a**).

**Figure 2:**
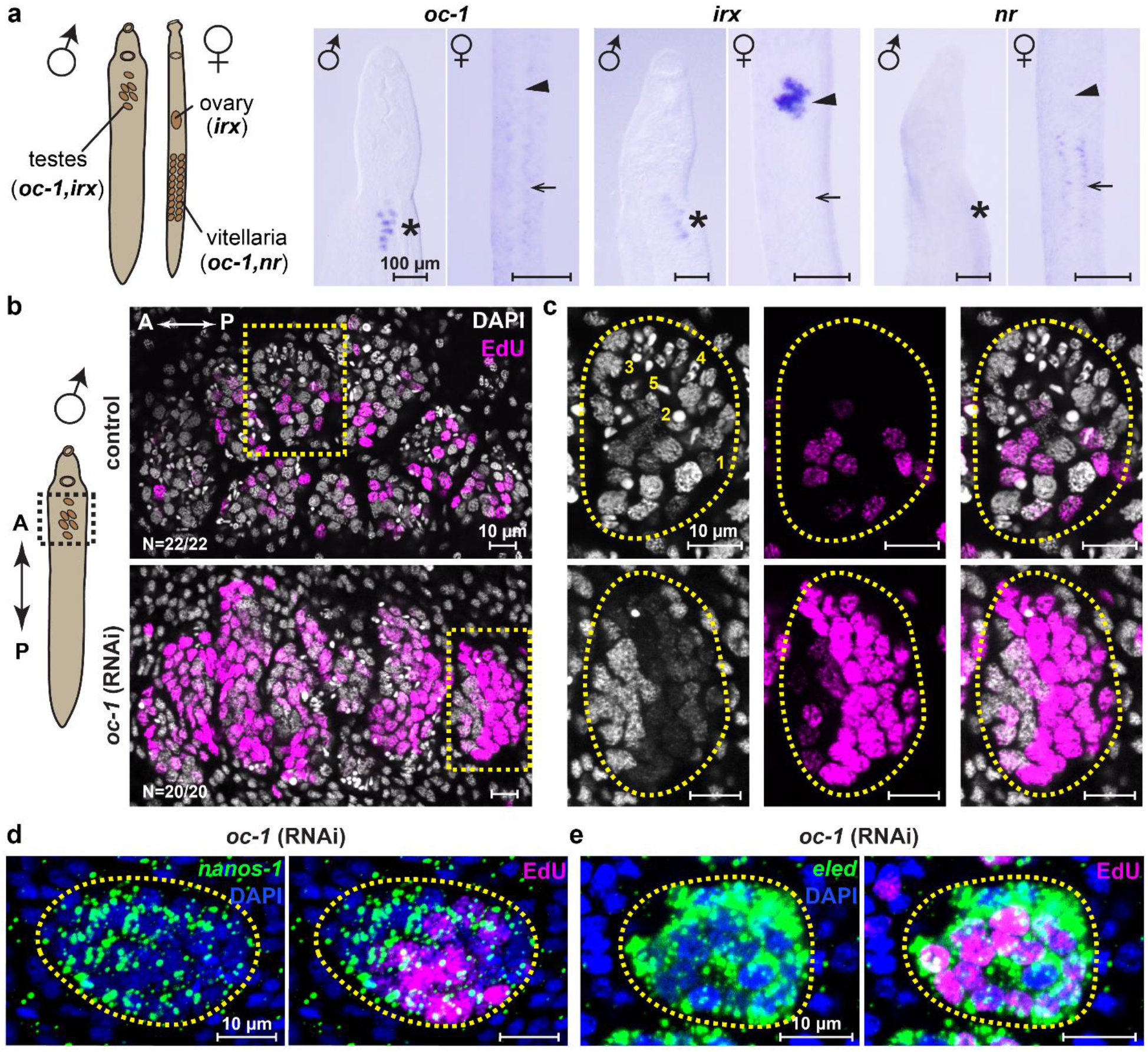
*oc-1* RNAi leads to over-proliferation of male GSCs. (a) WISH images of *oc-1, irx* and *nr* expression in male and female juvenile parasites. Left: schematic showing the locations of major reproductive organs in schistosome juveniles. Asterisks: primordial testes; arrowheads: ovary primordia; arrows: presumptive vitellaria primordia. (b) Confocal images of DAPI/EdU-stained testes in control and *oc-1* RNAi male juvenile parasites. The imaged area corresponds to the dashed box in the schematic on the left. A: anterior; P: posterior. N: number of parasites analyzed. (c) Magnified views of individual testis lobules corresponding to yellow boxes in (b). Dashed circles: testis lobe boundary. Nuclear morphologies are labeled in the image as (1) GSC (undifferentiated spermatogonium), (2) spermatocyte, (3) round spermatid, (4) elongating spermatid, and (5) sperm. Stage 2-5 are considered as differentiated germ cells. Note the increased fraction of EdU^+^ nuclei and reduced number of nuclei corresponding to differentiated germ cells after RNAi compared to controls. (d-e) FISH images showing broad expression of nanos-*1* (d) and *eled* (e) among GSCs, including EdU^+^ nuclei, after *oc-1* RNAi.

To evaluate the function of these TFs, we performed RNAi to knock them down in juvenile parasites. For these experiments, parasites were retrieved from mice 3.5 weeks post-infection and soaked in double-stranded RNA (dsRNA) continuously *in vitro* for 2 weeks. We focused on the male germline because female development is retarded under *in vitro* culture and therefore associated phenotypes are difficult to evaluate^5^. While *nr* or *irx* RNAi did not result in any noticeable phenotypes (**Supplementary Fig. 4**), *oc-1* RNAi led to a significant increase in the number of proliferating cells labeled by 5-Ethynyl-2’-deoxyuridine (EdU) and a dramatic reduction in the abundance of differentiated germ cells, including spermatocytes, spermatids and sperm (**Fig. 2b**). The EdU^+^ cells had a nuclear morphology of undifferentiated spermatogonia but were smaller in size compared to the nuclei in control samples, potentially due to the overcrowding of cells in individual testis lobules as a consequence of over-proliferation. Consistent with their spermatogonial identity, EdU^+^ cells express *nanos-1* and *eled* after *oc-1* RNAi (**Fig. 2d,e**), indicating that the expression of these GSC markers is independent of *oc-1* function. Together, these results suggest that *oc-1* is required not for maintaining the identity of GSCs but rather for suppressing the intrinsic tendency of proliferation in GSCs and promoting their differentiation.

### An RNAi screen identifies *boule* as a germline regulator that phenocopies *oc-1* after knockdown

To identify the pathway through which *oc-1* functions, we performed an RNAi screen to knock down genes enriched in GSCs identified by scRNAseq, focusing on predicted RNA binding proteins, enzymes, and receptors. The goal of these experiments was to identify genes that phenocopy *oc-1* upon RNAi. Out of the 39 genes knocked down (**Supplementary Table 2**), we observed male germline development defects in RNAi experiments of six genes (**Fig. 3a**). Of these, only knockdown of *boule* (Smp_144860) gave rise to a similar but stronger phenotype compared to that of *oc-1* RNAi (**Fig. 3b-e**): testis lobules contained densely packed EdU^+^ *nanos-1^+^/eled^+^* GSCs and almost completely lacked differentiated germ cells. Consistent with its phenotype, *boule* is co-expressed with *oc-1* in spermatogonial cells (**Supplementary Fig. 3**). *boule* is a member of the Deleted in Azoospermia (DAZ) family of RNA-binding proteins, which are known germline regulators with important roles in both GSC maintenance and gametogenesis, though specifics vary between animals^26–30^. To investigate whether other genes associated with DAZ function in other animals are also involved in schistosome germline development, we searched for potential DAZ-interacting partners, DAZAP and DZIP^27,47,48^, in the *S. mansoni* genome based on BLAST similarity. We only found one DZIP homolog, RNAi of which did not generate a germline phenotype (**Supplementary Fig. 5**), indicating that *boule* function in the schistosome is either independent of DZIP or involves other unidentified redundant interacting proteins.

**Figure 3:**
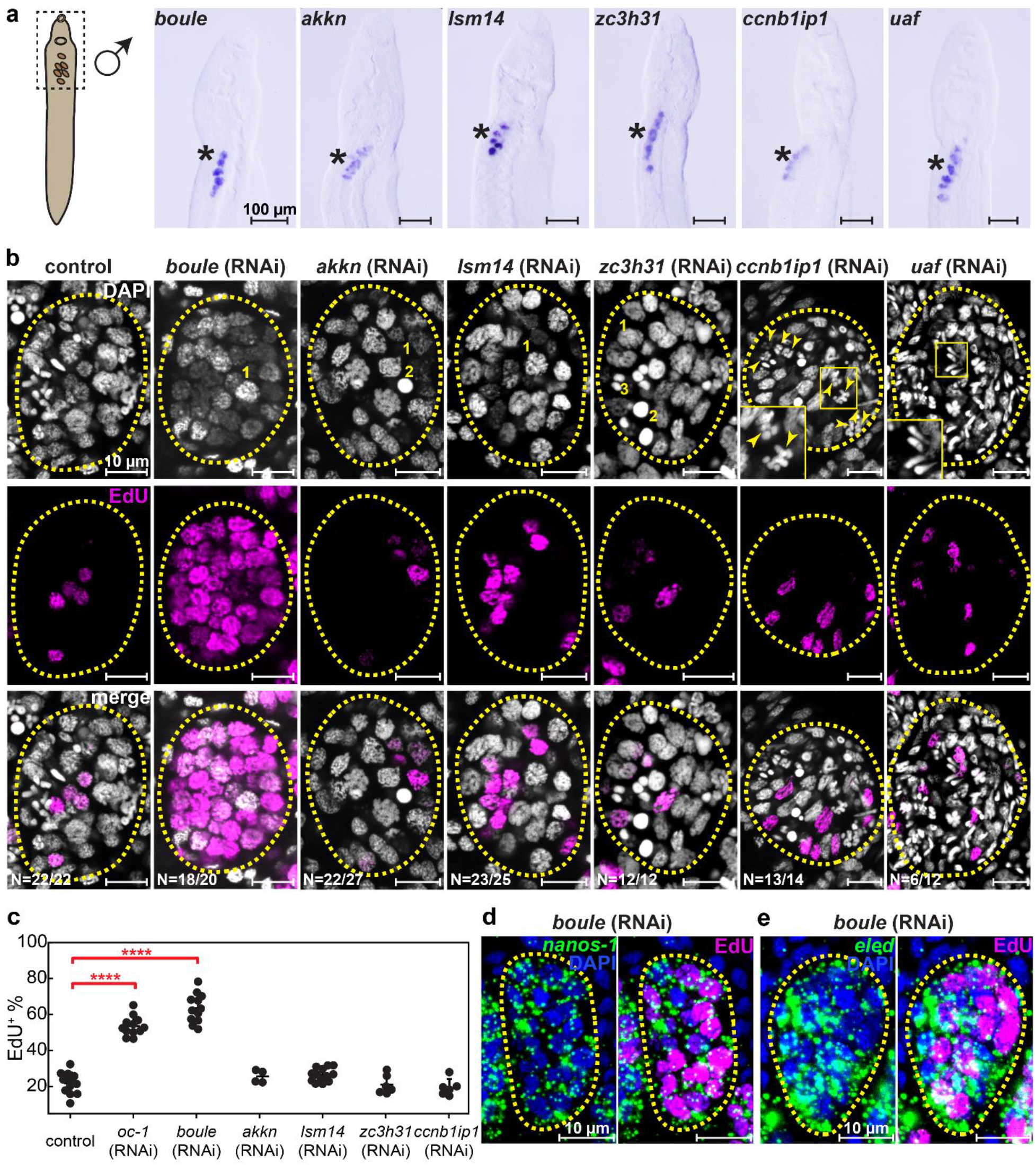
An RNAi screen identifies that *boule* phenocopies *oc-1* after RNAi in juvenile male germline. (a) WISH images showing the expression of *boule, akkn, lsm14, zc3h31, ccnb1ip1* and *uaf* in male juvenile parasites. Asterisks: testes. The image areas correspond to the dashed box in the schematic on the left. (b) Confocal images showing representative individual testis lobules stained by DAPI and EdU in control and *boule, akkn, lsm14, zc3h31, ccnb1ip1* and *uaf* RNAi juvenile parasites. Insets: magnified boxed areas. Dashed circles: testis lobe boundary. Arrowheads in *ccnb1ip1* RNAi image: incomplete separation of meiotic nuclei. N: number of parasites analyzed. Nuclear morphologies are labeled in the images as (1) GSC (undifferentiated spermatogonium), (2) spermatocyte, and (3) round spermatid. (c) Fraction of EdU^+^ nuclei in testes in control and RNAi parasites. Each data point represents the average across all testis lobules in a single parasite. Number of counted parasites: N = 12 (control); N = 12 (*oc-1* RNAi), N = 12 (*boule* RNAi), N = 4 (*akkn* RNAi); N = 13 (*lsm1* RNAi); N = 6 (*zc3h31* RNAi); N = 6 (*ccnb1ip1* RNAi). ****p ≤ 0.0001 (Student’s t-test). Error bars represent standard deviation. FISH images showing broad expression of nanos-*1* (d) and *eled* (e) in virtually all cells, including EdU^+^ cells, in testes after *boule* RNAi.

The other five genes identified in this screen affect germline differentiation but not proliferation. RNAi of a putative ankyrin repeat-containing protein kinase (*akkn*, Smp_131630) and a LSM14 domain-containing protein (*lsm14*, Smp_129960) blocked the differentiation of germ cells. RNAi of a zinc finger CCCH domain-containing protein (*zc3h31*, Smp_056280) and an E3 ubiquitin protein ligase CCNB1IP1 (*ccnb1ip1*, Smp_344490) caused defects in spermatogenesis. RNAi of a homolog of uro adherence factor A (*uaf*, Smp_105780) resulted in premature accumulation of sperm and abnormal GSC nuclear morphology (**Fig. 3b,c**).

Finally, to examine whether these genes play similar functional roles in homeostasis beyond development, we performed RNAi experiments on sexually mature parasites, retrieved from mice 5.5 weeks post-infection (**Fig. 4a**). **Fig. 4b,c** show that most phenotypes, including those induced by *oc-1* and *boule* knockdowns, are identical between juveniles and adults, except for *akkn*, which did not yield a knockdown phenotype in adult parasites (**Supplementary Fig. 6**), suggesting that its function is restricted to regulating development.

**Figure 4:**
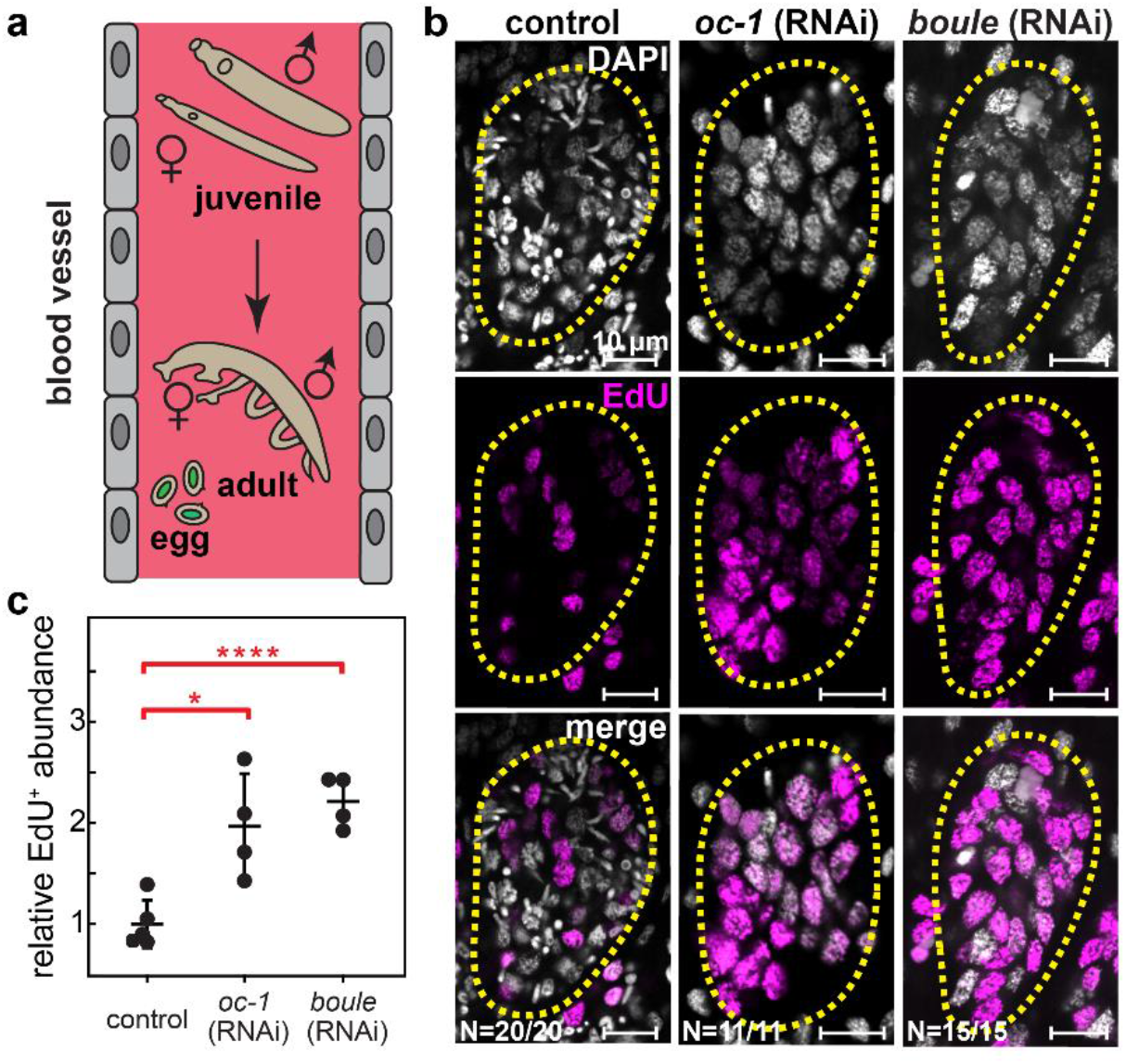
Knockdowns of *boule* and *oc-1* cause male GSC over-proliferation in adult parasites. (a) Schematic of sexual maturation process from juveniles to adults, which mate and lay eggs. (b) Confocal images showing representative individual testis lobules stained by DAPI and EdU in adult parasites after control, o*c-1* and *boule* RNAi. N: number of parasites analyzed. Dashed circles: testis lobe boundary. (c) Relative abundance of EdU^+^ nuclei per image area in testes normalized against the abundance in control parasites. Number of parasites counted: N = 5 (control); N = 4 (*oc-1* RNAi); N = 4 (*boule* RNAi). ****p ≤ 0.0001, *p ≤ 0.05, according to Student’s t-test. Error bars represent standard deviation.

### *oc-1* and *boule* expressions are mutually dependent and exhibit similar epistatic interactions with *nanos* and *eled*

The observation that RNAi phenotypes of *oc-1* and *boule* are mostly equivalent motivated us to investigate the nature of relationship between them. We followed up RNAi of either gene with WISH to evaluate the changes in the expression of both. We found that *boule* expression was undetectable after *oc-1* RNAi, and *oc-1* expression was also eliminated upon *boule* RNAi (**Fig. 5a**). This result suggests that *oc-1* and *boule* regulate (directly or indirectly) each other at the transcriptional level.

**Figure 5:**
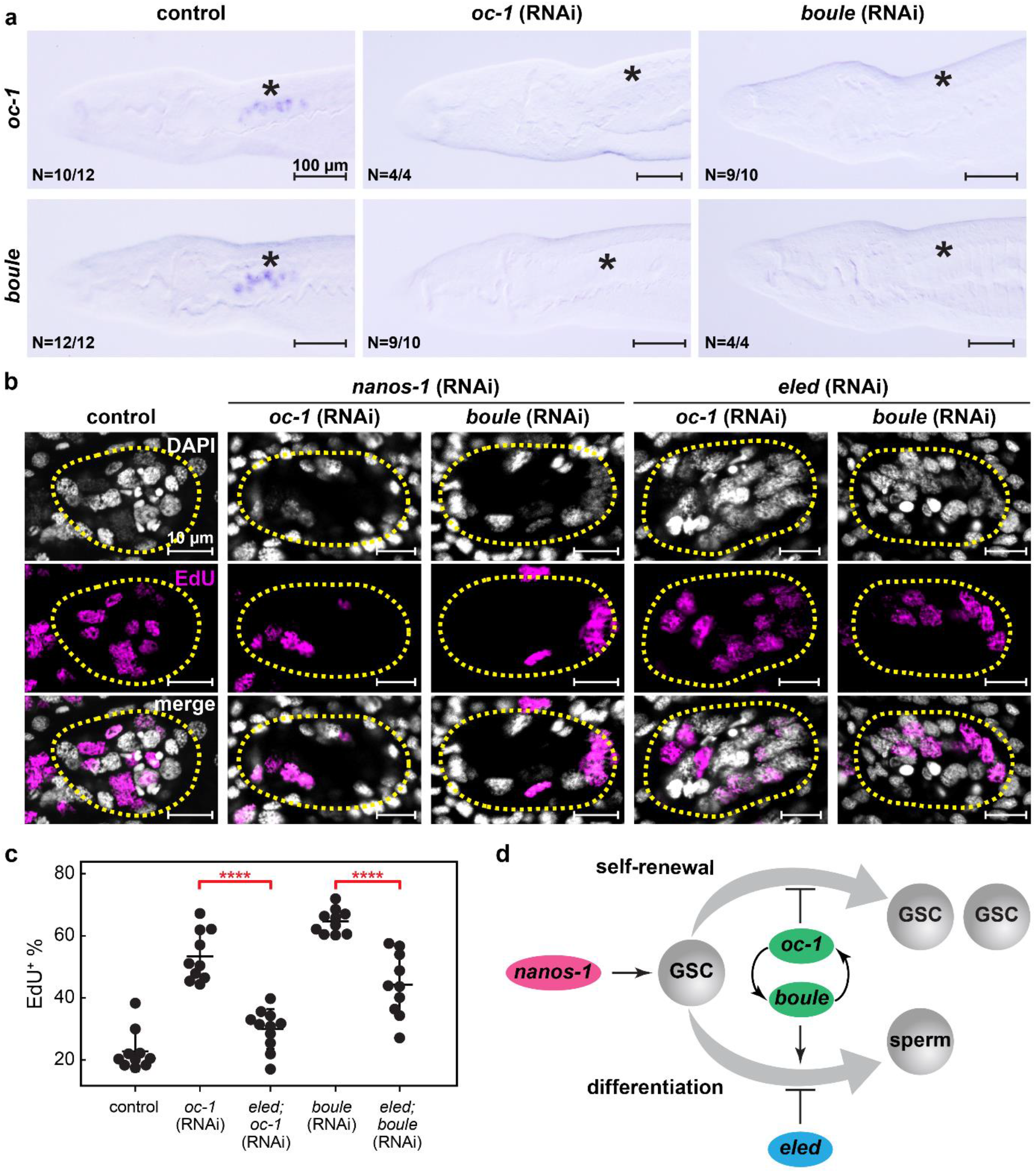
Epistatic interactions of *oc-1, boule, nanos-1* and *eled*. (a) WISH images showing that the expression of *oc-1* and *boule* are mutually dependent. Knockdown of either one causes the elimination of both. Asterisks: testes. N: number of parasites analyzed. (b) Confocal images showing representative individual testis lobules stained by DAPI and EdU in control and double knockdowns. Dashed circles: testis lobe boundary. (c) Fraction of EdU^+^ nuclei in testes in control and RNAi parasites. Each data point represents the average across all testis lobules in a single parasite. Number of parasites analyzed: N = 10 (control); N = 10 (*oc-1* RNAi); N = 11 (*eled;oc-1* double knockdown); N = 10 (*boule* RNAi); N = 10 (*eled;boule* RNAi). *nanos-1* RNAi experiments were not quantified because only a few GSC nuclei are present per testis lobe, which is insufficient for reliable statistics. ****p ≤ 0.0001, according to Student’s t-test. Error bars represent standard deviation. (d) Summary of epistatic interactions of *oc-1*, *boule*, *nanos-1* and *eled* in regulating male GSC proliferation and differentiation.

If *oc-1* and *boule* overlap through the same functional pathway, we expect them to exhibit identical epistatic interactions with other GSC regulators. Previously, we found that *nanos-1* is required for GSC proliferation and differentiation, knockdown of which causes degeneration of testes and loss of differentiated germ cells, whereas *eled* is thought to inhibit GSC differentiation, RNAi of which ectopically activates spermatogenesis and leads to premature accumulation of sperm^5^. As their RNAi phenotypes are distinct from those resulting from *oc-1* and *boule* RNAi, it is feasible to test epistatic interactions among this set of genes through double RNAi knockdowns.

Both *nanos-1;oc-1* and *nanos-1;boule* double knockdowns exhibited the phenotype of *nanos-1* single knockdown (**Fig. 5b**), suggesting that *nanos-1* likely functions upstream of *oc-1* and *boule* as a genetic suppressor. By contrast, *eled;oc-1* or *eled;boule* double knockdowns alleviated the defects observed in *oc-1* or *boule* single knockdowns, measured by the presence of differentiated germ cells at all spermatogenesis stages and the reduction in the number of EdU^+^ cells (**Fig. 5b,c**). These observations indicate that *oc-1*, *boule* and *eled* collectively maintain the balance between differentiation and proliferation of GSCs, with *oc-1* and *boule* suppressing proliferation and promoting differentiation versus *eled* suppressing differentiation (**Fig. 5d**). When the functions of both sides are lost after double knockdowns, the balance can be reestablished.

### Planarian homolog of *onecut* is also required for male germline development

To determine if the function of *onecut* in germline development is evolutionarily conserved, we studied the planarian flatworm *S. mediterranea*, a free-living cousin of schistosomes. Using BLAST sequence similarity search, we identified three planarian *onecut* homologs (**Supplementary Fig. 2**), but only one (*Smed-oc-1*, dd_Smes_v1_39638_1_1) is detected in the male germline (**Supplementary Fig. 7a,b**). Specifically, it is expressed in testis lobules, which are distributed beneath the dorsal epidermis of sexually mature planarians; in sexually immature planarians, *Smed-oc-1* expression is specific to testis primordia, which are essentially GSC clusters (**Fig. 6a**). High resolution images of individual testis lobules reveal that *Smed-oc-1* expression is restricted to spermatogonial stem cells and mitotic spermatogonia, but not detected in differentiated germ cells (**Fig. 6b**). Consistently, in asexual planarians, *Smed-oc-1* is also present in presumptive GSCs (**Supplementary Fig. 7c**).

**Figure 6:**
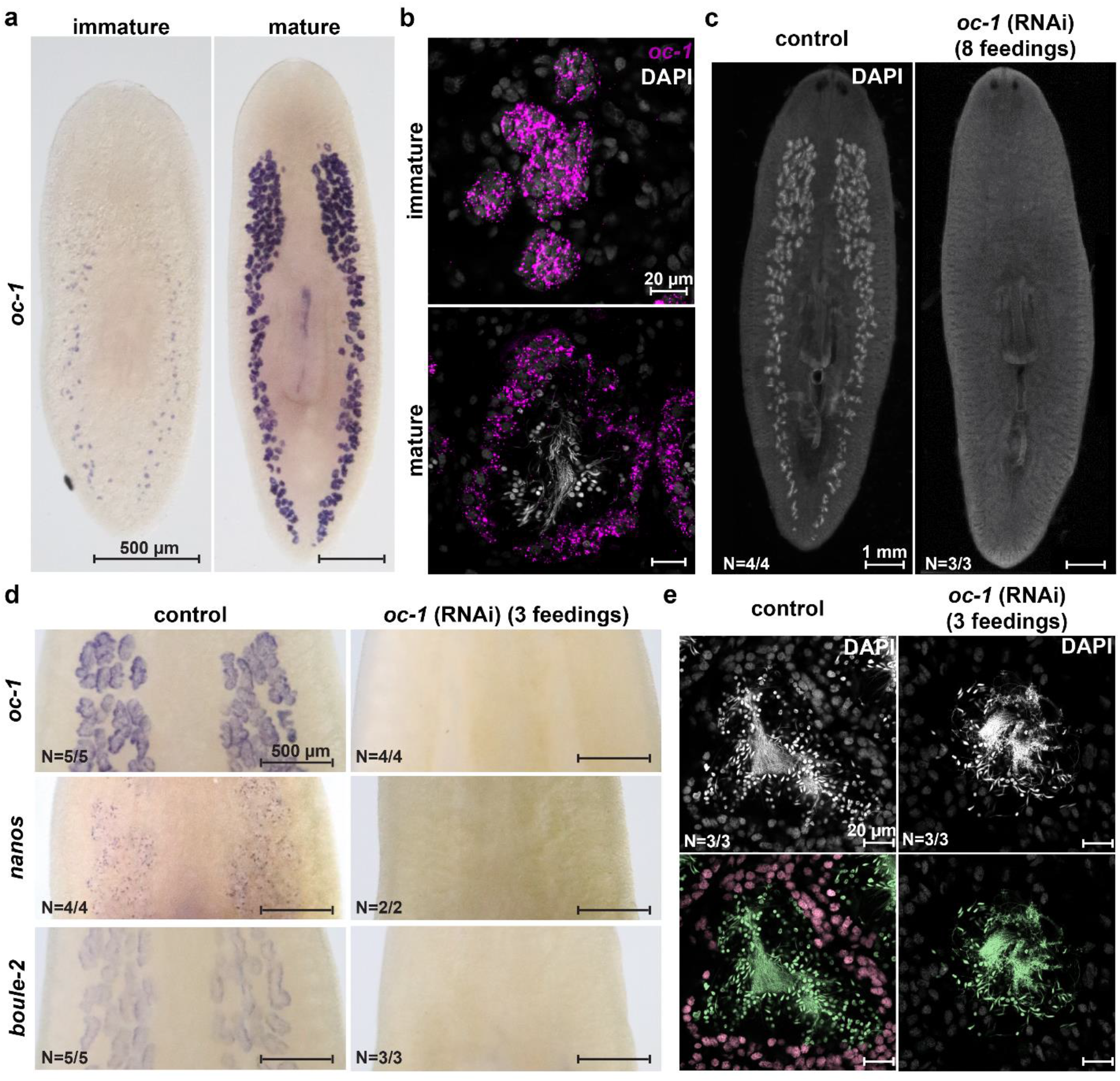
*Smed-oc-1* is required for the maintenance of spermatogonial cells. (a) WISH images showing the expression of *Smed-oc-1* in sexually immature (left) and mature (right) planarians. The punctuated signal in the immature planarian corresponds to presumptive testis primordia. In the mature planarian, the signal is specific to testes. (b) FISH of *Smed-oc-1* showing its expression in testis primordia within sexually immature planarians (top) and in spermatogonial cells (including GSCs and mitotic spermatogonia) in sexually mature planarians (bottom). (c) Dorsal view of sexually mature planarians after control and *Smed-oc-1* RNAi. Note that testes, which are distributed beneath the dorsal epithelium and visualized by strong DAPI signal, are completely eliminated after *Smed-oc-1* RNAi. Animals were fixed after eight feedings of dsRNA. N: number of worms analyzed. (d) WISH images showing that the expression of *Smed-oc-1, Smed-nanos*, and *Smed-boule-2* are disrupted after three feedings of *Smed-oc-1* RNAi. N: number of worms analyzed. (e) Single confocal sections of DAPI stained testes of control (left) and *Smed-oc-1* (right) RNAi planarians. Bottom: false colored images to annotate undifferentiated spermatogonial cells (pink) and differentiated germ cells (green). Animals were fixed after three feedings of dsRNA. Note that at this early time point of o*c-1* RNAi spermatogonial cells at the periphery are missing, while differentiated germ cells remain. N: number of worms analyzed.

Knockdown of *Smed-oc-1* led to the complete loss of testes (**Fig. 6c**), suggesting that it is required for maintaining the germline fate. This function is distinct from that of *Sm-oc-1*, which is dispensable for maintaining the identity of GSCs but necessary for regulating GSC proliferation and differentiation.

We next evaluated the relationship between *Smed-oc-1* and other known planarian germline regulators such as *nanos* and *boule*. The planarian *nanos* homolog, *Smed-nanos*, is a GSC marker and required for GSC specification and maintenance^17,24^. The planarian *boule* homologs, *Smed-boule-1* and *Smed-boule-2*, are also expressed in spermatogonial cells along with *Smed-oc-1*. *Smed-boule-1* is involved in spermatogenesis and loss of its function leads to the elimination of differentiated germ cells, whereas *Smed-boule-2* is necessary for the maintenance of GSCs (**Supplementary Fig. 7d**)^27^. We found that, at an early time point during *Smed-oc-1* RNAi, the expressions of *Smed-nanos* and *Smed-boule-2* were concomitantly abolished (**Fig. 6d**). This observation is consistent with the rapid depletion of spermatogonial cells, though differentiated germ cells were still present at this time point (**Fig. 6e**). These results suggest that *Smed-oc-1* function overlaps with that of *Smed-nanos* and *Smed-boule-2*, but not *Smed-boule-1*. Based on these observations, we conclude that the coexpression of *onecut, boule* and *nanos* in the male germline appears to be conserved between planarian and schistosome, but the functional relationship within this gene set has been rewired significantly.

## Discussion

Schistosome germline is essential for both disease transmission and pathology, as it produces eggs that serve as the transmissible agents and induce host pathological inflammatory response^13^. The germline develops post-embryonically at the juvenile stage from a population of stem cells with a somatic origin that is carried over from asexual larval parasites^5^. During the maturation of juvenile parasites, GSCs need to be regulated to proliferate and differentiate at rates such that somatic development can be completed before the production of gametes. In this work, we identify a germline specific stem cell regulatory program that balances the fate of GSCs between proliferation and differentiation. We also studied the evolutionary conservation and modification of this program by comparing the schistosome and its planarian cousin.

Our study uses scRNAseq to construct a complete map of the juvenile schistosome stem cell population at its intramammlian life-cycle stage. The stem cells are separated into nine populations, which are further divided into two groups based on their transcriptional signatures: the presumptive multipotent populations, including the GSCs, and the presumptive tissue specific progenitors that are of muscle, neural, intestinal/parenchymal, and epidermal lineages. Each progenitor population expresses a specific battery of genes associated with respective differentiated cell types, whereas multipotent populations exhibit a global repression of these differentiation genes. This stem cell subpopulation structure is strikingly similar to that of planarian stem cells (which contain pluripotent and lineage restricted progenitors^49–51^), raising the question of whether these subpopulations have one-to-one correspondence between the two animals. In this regard, our work establishes a model system and provides the data needed for comparative studies aiming at understanding stem cell evolution at the single-cell transcriptomic level.

The transcriptomic characterization of the schistosome GSCs led us to the discovery of a conserved regulatory gene set in the male germline, with *onecut* homeobox transcription factor and *boule* mRNA binding protein at its core. In the schistosome, *onecut* and *boule* expressions are mutually dependent; they form a genetic circuit that inhibits the over-proliferation of GSCs and promotes differentiation, balancing the effect of *eled*, which suppresses differentiation of GSCs. In the planarian, homologs of *onecut* and *boule* (specifically *Smed-oc-1* and *Smed-boule-2*) are instead required for GSC maintenance. Compared to the planarian, which has low fertility rate, the schistosome has impressive fecundity with each pair of sexually mature parasites being capable of laying hundreds of fertilized eggs per day^52,53^. Whether the regulatory modifications around *onecut* and *boule* are associated with the difference in reproductive output is an important question for future research. Our study lays the foundation by revealing the key members of this previously uncharacterized gene circuit that regulates GSC fate.

We expect the conservation of this germline specific regulatory program to extend beyond flatworms. *boule* homologs, along with other members of DAZ protein family, are known germline regulators in diverse animals from basal invertebrates to humans^26–30^, though their premeiotic functions in GSCs as reported here are considered to be rare among invertebrates^27^. By contrast, *onecut* homologs have been studied in the context of neural, liver, and pancreatic development using different animal models^32–37^, but its function in germline development and its interaction with *boule* were previously unknown. Of interest to our study is the observation that *onecut* homologs are indeed expressed in rodent and human testes^31–34^, suggesting that its germline function may be more universal than previously acknowledged.

An important question raised by our results is what factors maintain the identity of GSCs. Our previous work suggests that GSCs are specified from *eled*^+^ stem cells during the early juvenile development by turning on *nanos-1*^5^. In the current scRNAseq dataset, we noticed that GSCs are distinguished from other stem cell populations in the presumptive multipotent group only by a small number of genes, while sharing a large set of common transcriptional signatures. The GSC-specific genes only contain three TFs, but none of them are required for maintaining GSC identity. Knockdown of other genes enriched in GSCs mostly caused defects in differentiation and spermatogenesis but did not affect the identity of GSCs either. These observations favor the hypothesis that GSC identity may be maintained by extrinsic cues, though regulation by other intrinsic determinants such as post-transcriptional regulatory factors cannot be excluded.

In mammalian testes, these extrinsic cues are provided by discrete somatic niche cells (e.g., Sertoli cells) to support male GSCs^54,55^; in *Drosophila* and *C. elegans*, cap/hub cells form the niche instructing GSC fates through several signaling pathways^56,57^; in the planarian, somatic gonadal cells required for male GSC specification and maintenance have also been defined^58,59^. All these “niche” cells are in close spatial proximity with their GSC counterparts. However, in the schistosome testes, most GSCs are not in contact with any somatic tissues. This is particularly clear in *oc-1* RNAi or *boule* RNAi animals where the testis lobules contain exclusively a large number of densely packed cycling GSCs. These observations suggest that the extrinsic cues for maintaining GSC identity in schistosomes is likely systemic, potentially through neural or hormonal regulations. This hypothesis is particularly compelling because schistosome maturation is known to depend on host immune cues^12,13^. The host signals need to be sensed by parasite cells at the host-parasite interface, then translated and relayed internally to instruct GSC activities^60^. In the planarian, germline development is regulated by neuroendocrine signals, such as neuropeptides^61^, implicating that there may be a similar crosstalk between the nervous system and the gonads in schistosomes. Here our characterization of the intrinsic regulatory program of the schistosome GSCs opens the door for exploring these important questions. Given the significance of parasite germline in disease transmission and pathology, better understanding of the basic biology of this developmental process will help to define vulnerable points for combating this major parasitic disease.

## Methods

### Animals

*S. mansoni* juveniles and adults were retrieved from infected female Swiss Webster mice (NR-21963) by hepatic portal vein perfusion with 37°C DMEM supplemented with 5% heat inactivated FBS at 3.5 weeks and 5.5 weeks post-infection, respectively. Worms were cultured at 37°C in Basch Media 169 supplemented with 1X Antibiotic-Antimycotic^11^. In adherence to the Animal Welfare Act and the Public Health Service Policy on Humane Care and Use of Laboratory Animals, all experiments with and care of mice were performed in accordance with protocols approved by the Institutional Animal Care and Use Committees (IACUC) of Stanford University (protocol approval number 30366).

Sexual and asexual *S. mediterranea* were maintained in the dark at 18°C in 0.75× Montjuïc salts or 0.5 g/L Instant Ocean Sea Salts supplemented with 0.1 g/L sodium bicarbonate, respectively. Planarians were fed calf liver paste once or twice weekly and starved at least 5 d prior to fixation. For early knockdown experiments, large sexual planarians were chosen to ensure sexual maturity. Smaller animals (<4 mm in length) were used to test expression in sexually immature animals.

### scRNAseq and data analysis

Single cell suspensions of *S. mansoni* at juvenile stage were collected as previously described^5,25^ with several modifications. Specifically, juvenile parasites were transferred into 15 mL conical tubes and rinsed twice with PBS pre-warmed at 37°C. The parasites were dissociated in 4 mL of 0.25% trypsin diluted in HBSS for 20 min. Cell suspensions were passed through a 100 μm nylon mesh (Falcon Cell Strainer) and immediately doused with 11 mL of PBS supplemented with 1% BSA. The sample was then centrifuged at 300 g for 5 min at 4°C. Pellets were gently resuspended in PBS with 0.5% BSA, passed through a 30 μm nylon mesh, and stained with 5 μg/mL DAPI for 30 min. Stained samples were then washed and resuspended in PBS with 0.5% BSA and loaded on a SONY SH800S cell sorter. Dead cells were excluded based on DAPI fluorescence. Droplets containing single cells were gated using forward scattering (FSC) and side scattering (SSC). Cells that passed these gates were sorted into 384-well lysis plates containing Triton X-100, ERCC standards, oligo-dT, dNTP, and RNase inhibitor.

Reverse transcription and cDNA pre-amplification were processed with Smart-seq2 protocol as previously described^25^. We performed 23 cycles of PCR amplification and diluted the resulting cDNA in Qiagen EB buffer at 1:20 dilution as we empirically determined to yield an average final concentration of 0.4 ng/μL. Diluted cDNA were then tagmented and barcoded using in-house Tn5 tagmentase and custom barcodes fitted for 384-well plate. Library fragments concentration and purity were quantified by Agilent bioanalyzer and qPCR. 2 × 150 bp paired-end sequencing was performed on a NovaSeq 6000 at ~500,000 reads depth per cell at the Chan Zuckerberg Biohub Genomics core.

Raw sequencing reads were demultiplexed and converted to fastq files using bcl2fastq. Paired-end reads were mapped to *S. mansoni* reference genome version WBPS13 (WormBase Parasite) using Salmon (version 0.14.0) with “--validateMappings” flag. We performed downstream preprocessing and analysis on the estimated read counts in the Salmon output. Reads were initially mapped to exonic regions of the genome. To calculate total gene counts, read counts from individual exons of each gene were summed together. To eliminate low-quality cells, we filtered out cells with fewer than 520 genes detected, passing 7,657 cells for downstream analysis. This cut-off was chosen to bisect the bimodal distribution of the number of genes detected per cell.

Raw gene counts were normalized such that each cell has a total number of counts equal to the median library size of all cells. The resulting data were added with a pseudocount of 1 and Log2-transformed. Log2-transformed gene expressions with values less than 1 were set to zero, and genes detected in less than 1% or greater than 99% of cells were filtered out. The SAM algorithm was run with parameters ‘weight_PCs=False’ and ‘preprocessing=’StandardScaler’’^25^. SAM outputs 2D UMAP projections that we use for visualization.

Sub-clustering the stem cell populations was done by running SAM on *ago2-1*^+^ cells. We manually determined cluster identity based on marker gene expressions. We computed the centroid of each cluster in the principal component (PC) space generated by SAM. We applied hierarchical clustering using the average linkage method and the correlation distance metric to these coordinates to generate the dendrogram shown in **Fig. 1c**.

For each cluster, the marker genes were identified by computing partial sums of gene dispersions from the *k*-nearest-neighbor-averaged data:

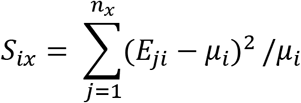

where *E_ji_* is the nearest neighbor averaged expression value for cell *j* and gene *i*, *μ_i_* is the mean expression of gene *i*, *n_x_* is the set of cells in cluster *x*, and *S_ix_* is the unnormalized maker score for gene *i* in cluster *x*. To account for differences across clusters, we normalize the marker score by the maximum value, 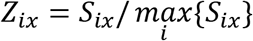. We rank the genes based on their marker scores, and only keep those that have greater than 0.5 SAM weight, greater than 0.1 normalized marker score, and exhibit >2 fold enrichment with respect to cells outside the cluster.

### RNAi

RNAi was performed using previously described protocols^5,6^. Clones were generated using oligonucleotide primers listed in **Supplementary Table 2-3** to amplify the gene fragment from cDNA and cloned into the vector pJC53.2 (Addgene Plasmid ID: 26536)^60^. To knock down genes in the schistosome, 10-15 juvenile or 6-8 adult parasites were soaked in Basch media 169 supplemented with ~20 μg/mL dsRNA for 2 weeks. The media with dsRNA were changed daily. Each RNAi was performed with at least two biological replicates. For gene knockdown in planarians, dsRNA was mixed with liver paste at a concentration of 70 ng/μL dsRNA and fed to planarians every 4 days for 3-8 feedings. dsRNA matching the *ccdB* and *camR*-containing insert of pJC53.2 was used as a control.

### In situ hybridization

RNA WISH and FISH experiments were performed following the established protocol^5,25,62^. Briefly, schistosomes were killed in 6 M MgCl_2_ for 30 s, fixed in 4% formaldehyde supplemented with 0.2% Triton X-100 and 1% NP-40 for 4 hr, and then dehydrated in methanol. Adult parasites were relaxed by 0.25% ethyl 3-aminobenzoate methanesulfonate for ~1 min to separate male and female parasites before fixation. Dehydrated juveniles were bleached in 3% H_2_O_2_ for 30 min, rehydrated, permeabilized by 10 μg/mL proteinase K for 20 min, and then post fixed with 4% formaldehyde. Dehydrated adults were rehydrated, bleached in the bleaching solution (5% formamide, 0.5× SSC, 1.2% H_2_O_2_) for 45 min under bright light, permeabilized by 10 μg/mL proteinase K for 30 min, and then post fixed with 4% formaldehyde. The hybridization was performed at 52°C with riboprobes, which were then detected through either alkaline phosphatase-catalyzed NBT/BCIP reaction (for WISH) or peroxidase-based tyramide signal amplification (for FISH). Planarians were killed with ice-chilled 5% NAC solution for 10 min and then fixed for 2 hr in 4% formaldehyde with 1% NP-40, then dehydrated in methanol. Samples were then rehydrated, bleached for 2 hr in the bleaching solution under bright light, permeabilized with 10 μg/mL proteinase K for 10 min and post fixed with 4% formaldehyde. The rest of the steps were performed similarly to those described for schistosome in situ hybridization experiments, except that the hybridization was performed at 56°C. To compare gene expression between control and RNAi conditions using WISH, the development of WISH signal was progressed in parallel and simultaneously stopped.

### EdU labeling

EdU (TCI Chemicals) was dissolved in DMSO at 10 mM and added into medium at a final concentration of 10 μM. Both juvenile and adult schistosomes were pulsed with EdU overnight. Parasites were fixed and permeabilized as in *in situ* hybridization experiments. EdU incorporation was detected by click reaction with 25μM of Cy5-azide conjugates (Click Chemistry Tools) or Carboxyrhodamine 110 Azide conjugates (Click Chemistry Tools)^25^. To combine EdU detection with FISH, click reaction was performed after tyramide signal amplification.

### Imaging

Samples for fluorescence imaging were mounted in the scale solution (30% glycerol, 0.1% Triton X-100, 2 mg/mL sodium ascorbate, 4 M urea in PBS)^63^ and imaged on a Zeiss LSM 800 confocal microscope using either a 20× (N.A. = 1.0, working distance = 1.8 mm) water-immersion objective (W Plan-Apochromat) or a 40× (N.A. = 1.1, working distance = 0.62 mm) water-immersion objective (LD C-Apochromat Corr M27). To obtain confocal images of schistosome testes, the midplane of testis lobules was first determined, then ~7 confocal sections with optimal z spacing recommended by Zen software were recorded around the midplane to generate maximum intensity projections. To generate false color images (**Fig. 6e**), nuclei were annotated manually and recolored on top of the original DAPI images using a customized MATLAB script. WISH samples were mounted in 80% glycerol with 10 mM Tris supplemented with 1 mM EDTA, pH = 7.5 for imaging.

### Data and code availability

The schistosome stem cell scRNAseq data generated in this study is available through the Gene Expression Omnibus (GEO) under accession number GSE147355. The SAM analysis source code can be found at https://github.com/atarashansky/self-assembling-manifold.

## Acknowledgements

*S. mansoni* (strain: NMRI) was provided by the NIAID Schistosomiasis Resource Center for distribution through BEI Resources, NIH-NIAID Contract HHSN272201000005I. We thank M. Khariton, Y. Fan, S. Korullu and R. Jones for experimental help. Sequencing was performed at the Biohub. Y.X. and A.J.T. are supported by the Stanford Bio-X Interdisciplinary Graduate Fellowship. B.W. is supported by a Beckman Young Investigator Award.

## Author contributions

P.L. and B.W. designed the research, Y.X. and P.L. performed the cell sorting and scRNAseq experiments, A.J.T. and P.L. analyzed the scRNAseq data, P.L., D.N.S., and X.Y. performed the functional experiments, P.L. and B.W. wrote the paper with input from all other authors, S.R.Q. and B.W. supervised the project and provided conceptual advice.

## Supplementary Information for

**Supplementary Fig. 1:**
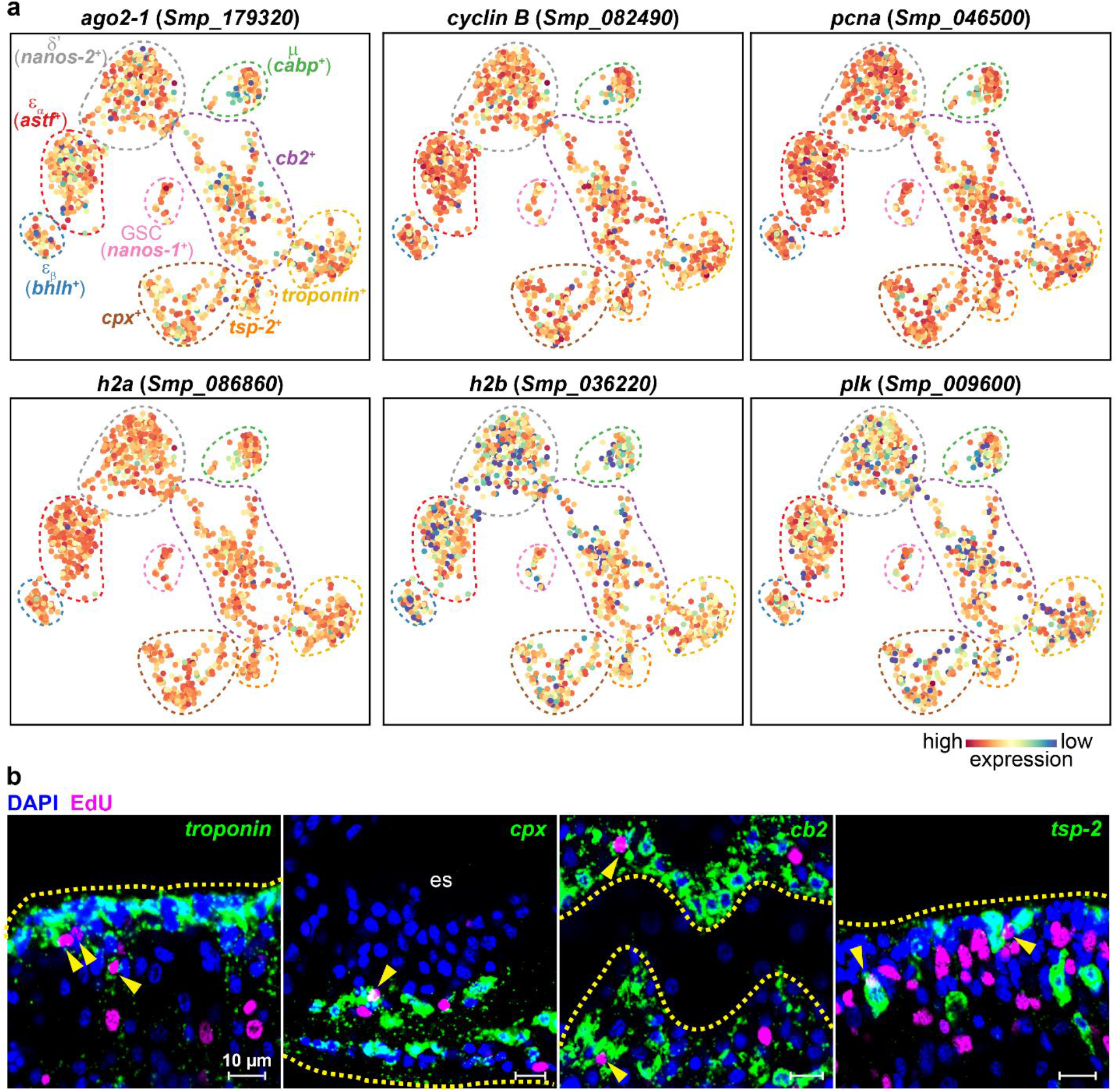
Characterization of presumptive progenitor populations. (a) UMAP projections showing that stem cell markers are ubiquitously expressed at high levels among *ago2-1^+^* cells, including all the progenitor populations. (b) EdU^+^ cells are detected within *troponin^+^, cpx^+^, cb2^+^* and *tsp-2^+^/Sm25^+^* populations, with examples highlighted by arrowheads. The marker genes used for FISH are *troponin* (Smp_018250)*, cpx* (Smp_050220)*, cb2* (Smp_141610) and *tsp-2* (Smp_335630). Dashed outline: parasite surface in *troponin, cpx*, and *tsp-2* FISH images and intestinal branches in *cb2* FISH image. es: esophagus.

**Supplementary Fig. 2:**
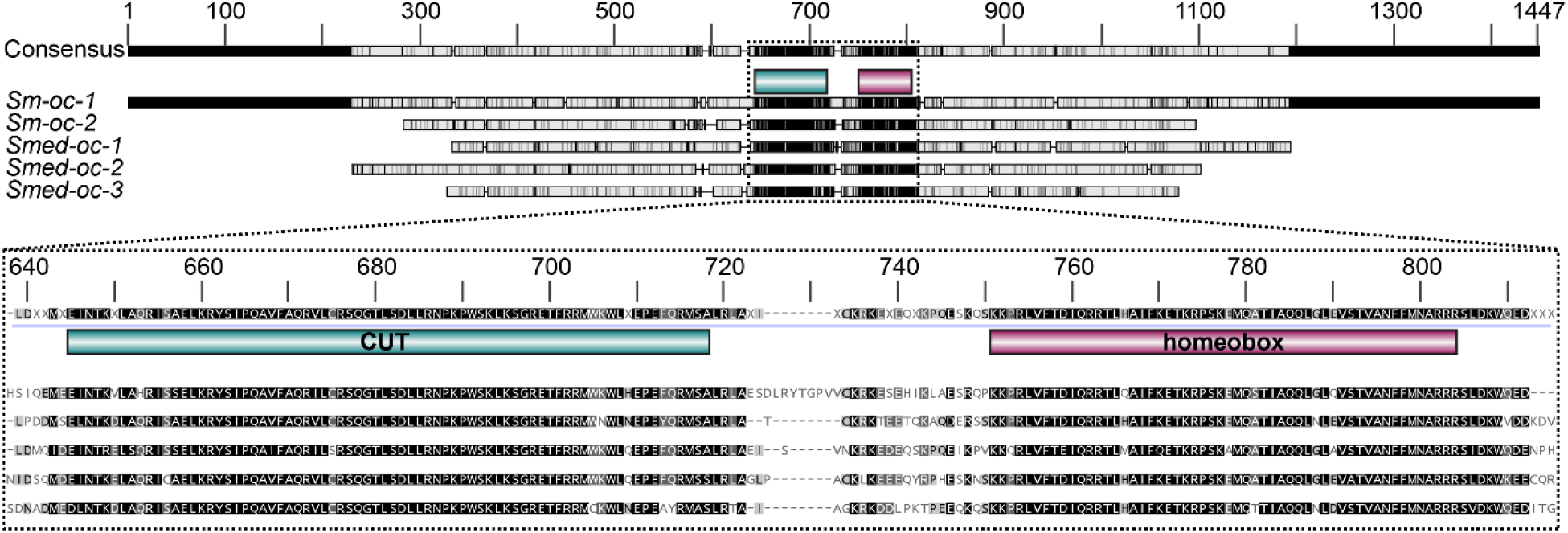
Sequence alignment of *onecut* transcription factors identified in the *S. mansoni* and *S. mediterranea* genomes. The alignment is performed using the multiple align function of Geneious prime with default settings. *onecut* homologs contain a CUT domain (green) and a homeobox domain (magenta). The domains are annotated by the InterProScan with the application of PfamA on the consensus sequence. Grey scale denotes the similarity of amino acids. Black: 100% similarity; white: less than 60% similarity. *Sm-oc-2* (Smp_013070) is not detectable in juvenile schistosomes by in situ hybridization. The gene IDs (PlanMine) of planarian *onecut* homologs are dd_Smes_v1_39638_1_1 (*Smed-oc-1*), dd_Smes_v1_39585_1_2 (*Smed-oc-2*), and dd_Smes_v1_35775_2_1 (*Smed-oc-3*).

**Supplementary Fig. 3:**
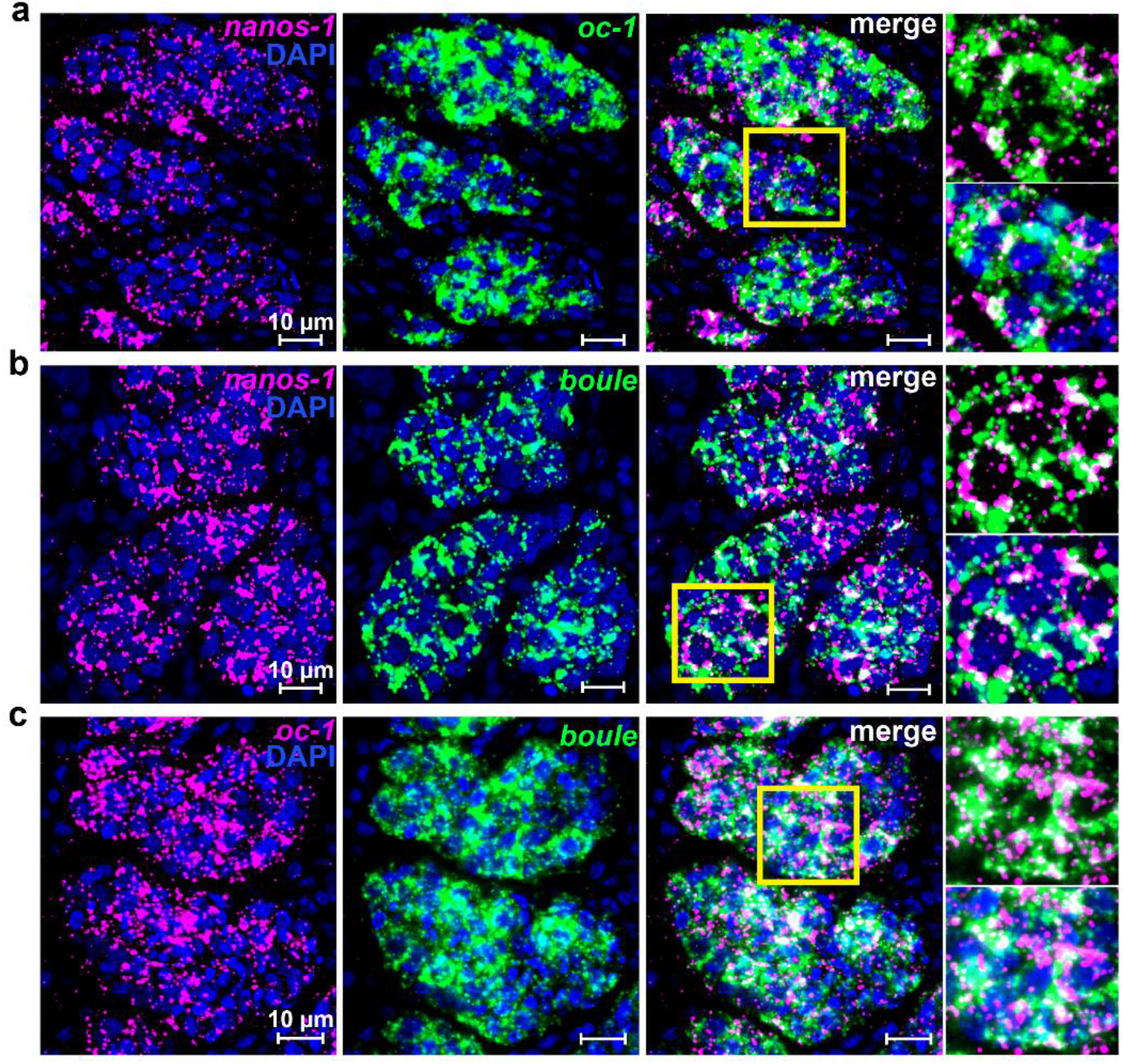
*Sm-oc-1* and *Sm-boule* are coexpressed in a subset of *nanos-1^+^* cells. Double FISH of (a) *nanos-1* and *oc-1*, (b) *nanos-1* and *boule* showing that not all *nanos-1^+^* cells express *oc-1* and *boule*, but all *oc-1^+^* and *boule^+^* cells express *nanos-1*. (c) Double FISH of *oc-1* and *boule* showing their colocalization. These expression patterns are consistent with scRNAseq results. Experiments were performed on juvenile parasites. Magnified views of boxed areas are shown to the right of the merge panel.

**Supplementary Fig. 4:**
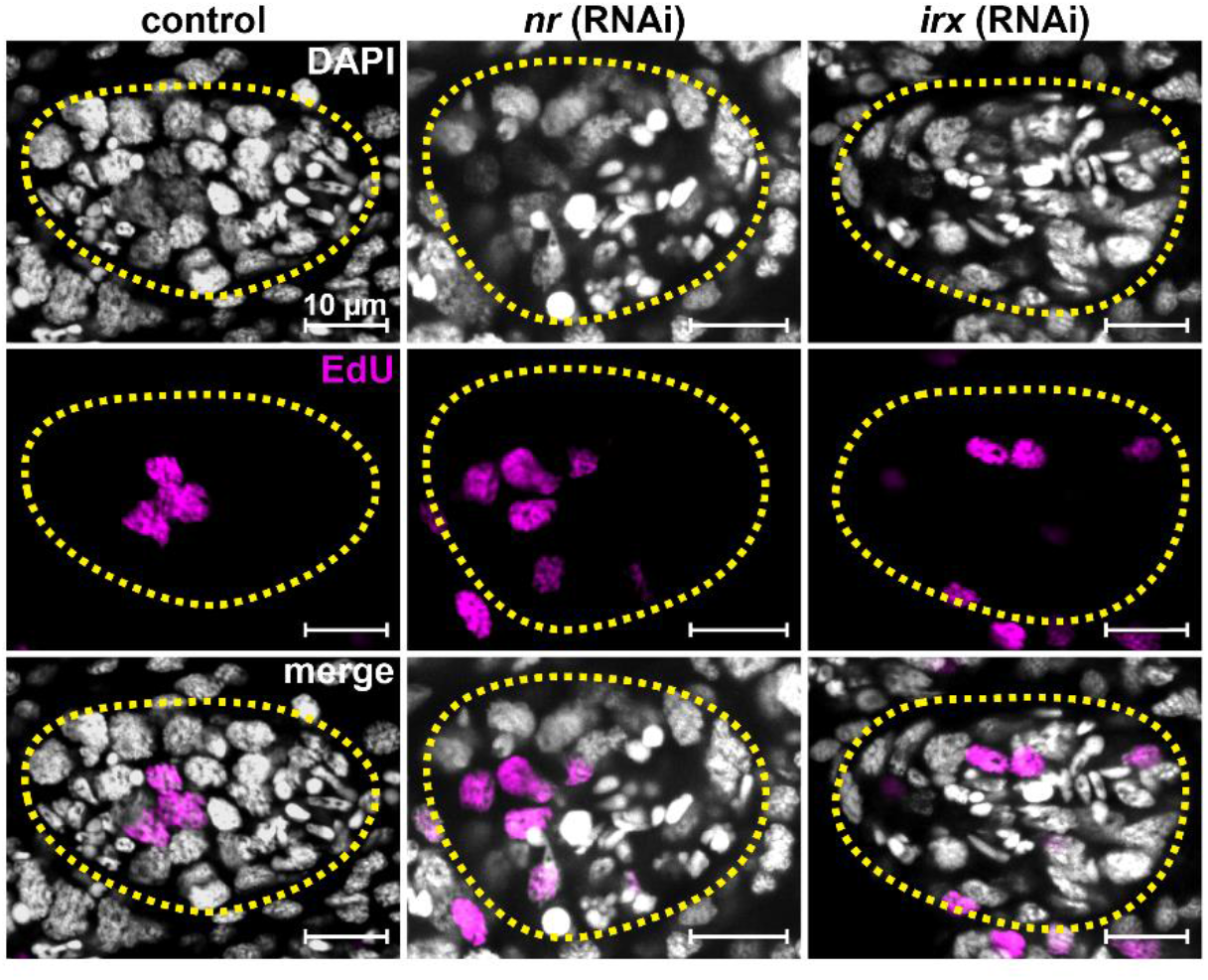
RNAi of *irx* and *nr* did not cause any noticeable phenotype. Confocal images of representative individual testis lobules stained with DAPI and EdU after control, *irx* RNAi and *nr* RNAi in juveniles. Dashed circles: testis lobe boundary.

**Supplementary Fig. 5:**
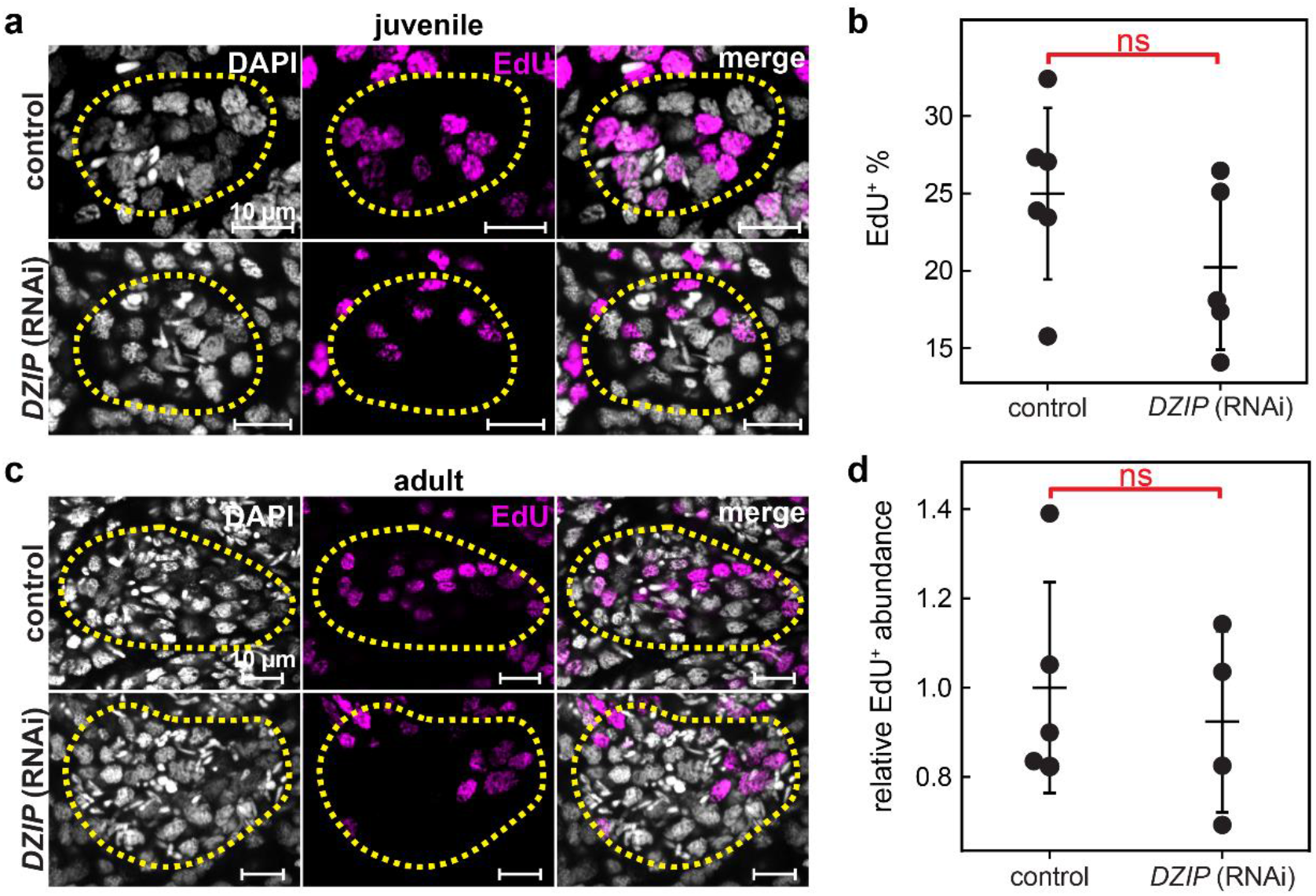
*DZIP* RNAi did not cause any noticeable phenotype in either juvenile or adult parasites. (a) Confocal images showing representative individual testis lobules to compare control and *DZIP* RNAi in juveniles, with the fraction of EdU^+^ nuclei quantified in (b). Number of parasites analyzed: N = 6 (control); N = 5 (*DZIP* RNAi). Data on adult parasites are reported in (c) with the relative abundance of EdU^+^ nuclei per image area normalized against the abundance in control parasites shown in (d). Number of parasites analyzed: N = 5 (control); N = 4 (*DZIP* RNAi). Dashed circles: testis lobe boundary. In (b,d), ns: no statistical significance according to Student’s t-test. Error bars represent standard deviation.

**Supplementary Fig. 6:**
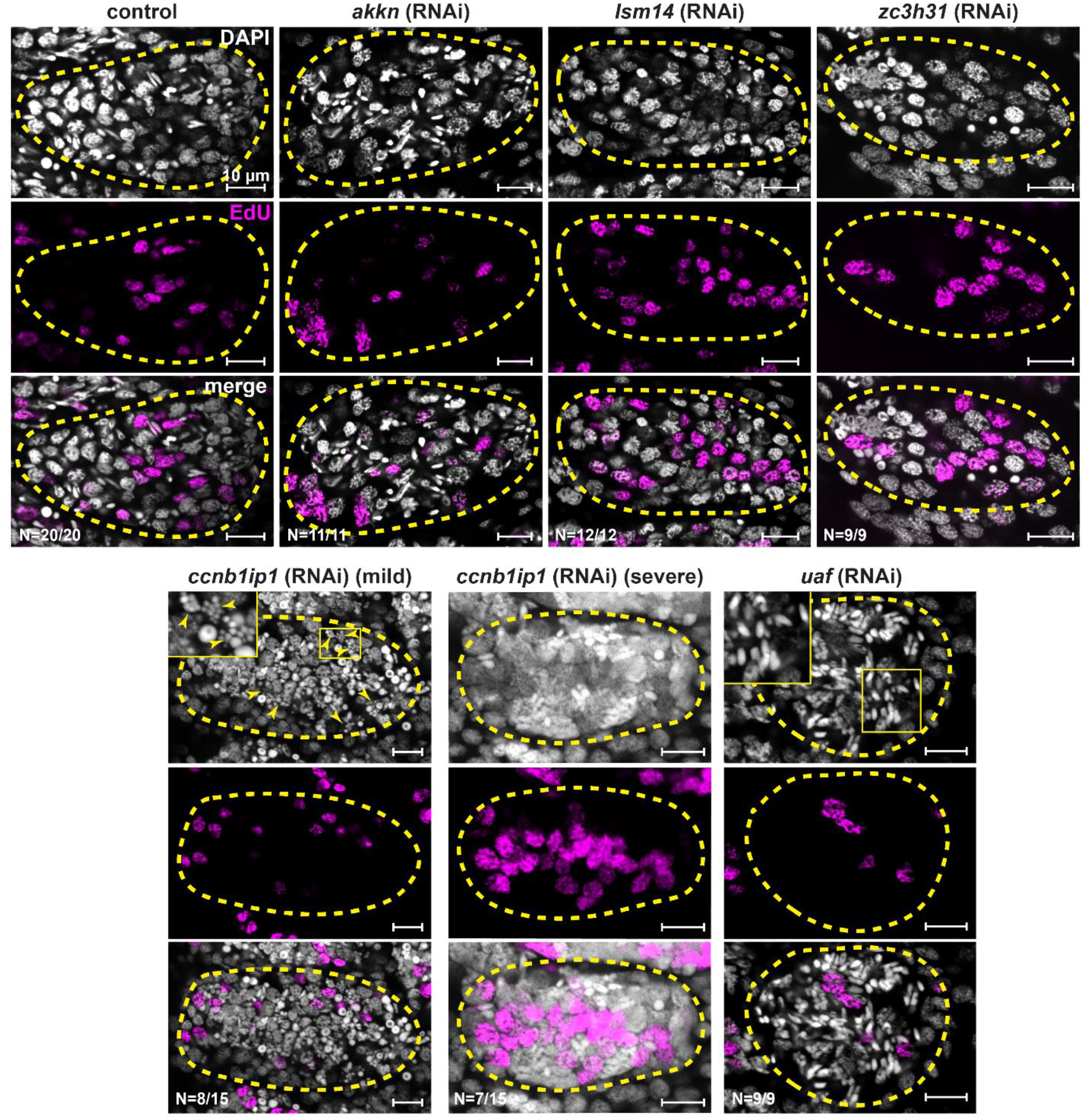
RNAi phenotypes of GSC regulators in adult parasites. Confocal images showing representative individual testis lobules stained by DAPI and EdU in control and *akkn, lsm14, zc3h31, ccnb1ip1* and *uaf* RNAi. *ccnb1ip1* RNAi can result in both a mild phenotype in which the separation of meiotic nuclei is incomplete (arrowheads), and a severe phenotype, in which nuclei appear to fuse between many cells. Insets: magnified boxed areas. Dashed circles: testis lobe boundary. N: number of parasites analyzed.

**Supplementary Fig. 7:**
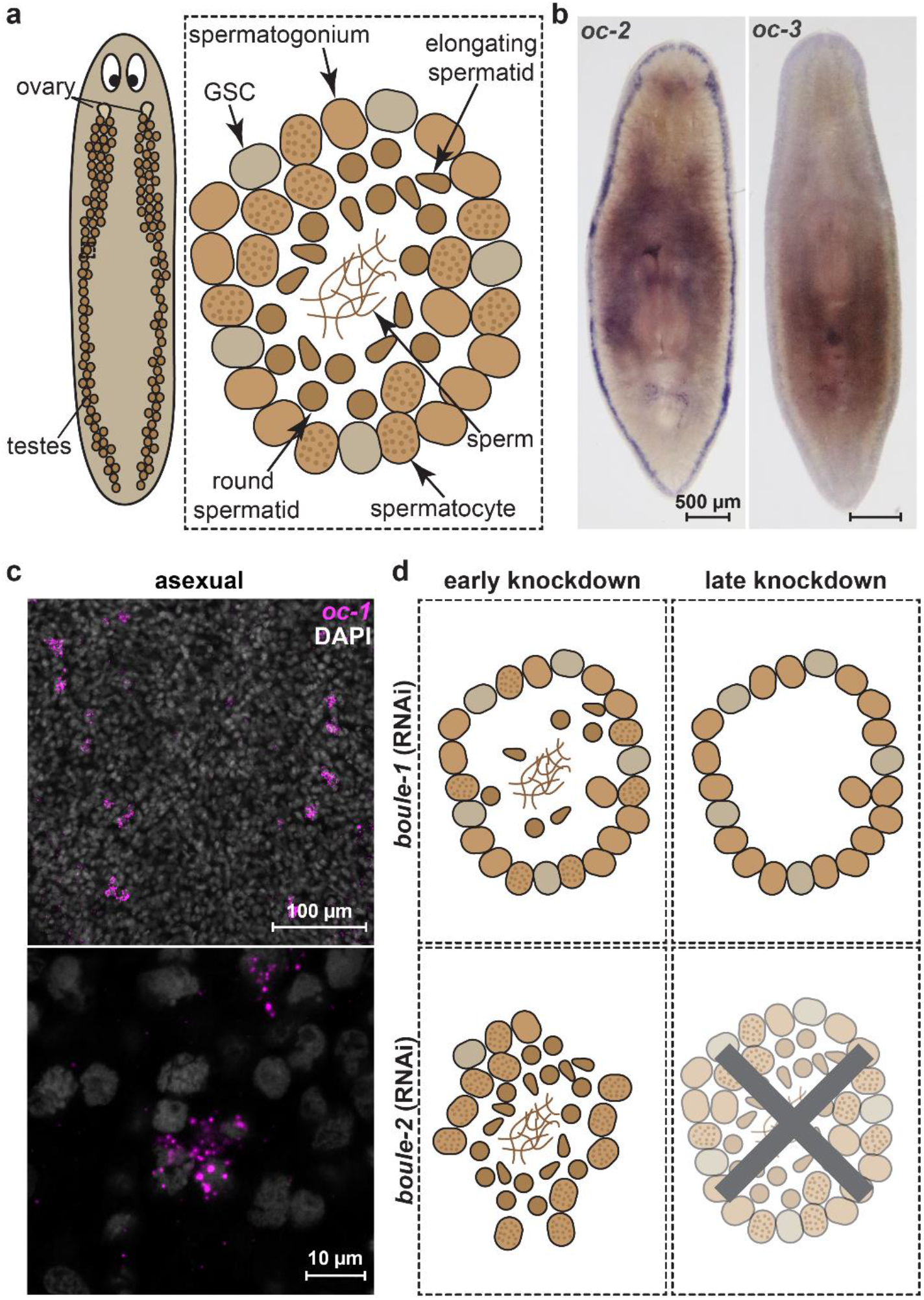
Expression of planarian *onecut* homologs. (a) Schematic of planarian testes lobules containing spermatogonial cells (GSCs and mitotic spermatogonia) at the periphery and differentiated germ cells (spermatocytes, round spermatids, elongating spermatids, and sperm) distributed in the internal layers. As in the schistosome, these cell types can be distinguished based on nuclear morphology after DAPI staining. (b) WISH images showing the expression of *Smed-oc-2* and *Smed-oc-3* in sexually mature planarians. (c) FISH images showing that *Smed-oc-1* is also detected in GSC clusters in asexual planarians. (d) Schematic showing the phenotypes observed after *Smed-boule-1* RNAi and *Smed-boule-2* RNAi as reported in ref. 27. After *Smed-boule-1* RNAi, differentiated germ cells are eliminated, whereas *Smed-boule-2* RNAi causes rapid loss of spermatogonial cells and eventually full degeneration of the testes.

